# The impacts of contemporary logging after 250 years of deforestation and degradation on forest-dependent threatened species

**DOI:** 10.1101/2023.02.22.529603

**Authors:** Michelle Ward, Kita Ashman, David Lindenmayer, Sarah Legge, Gareth Kindler, Timothy Cadman, Rachel Fletcher, Nick Whiterod, Mark Lintermans, Philip Zylstra, Romola Stewart, Hannah Thomas, Stuart Blanch, James E.M. Watson

## Abstract

Despite the importance of safeguarding forests and woodlands for achieving global climate and biodiversity agendas, logging continues across most forested countries. Forestry advocates often claim logging has minimal impacts, but rarely consider the cumulative threat deforestation and degradation has had, and continue to have, on species. Using New South Wales (Australia) as a case study, we quantify the extent of deforestation and degradation from 1750 – current. Using these estimates of overall loss as a baseline, we then quantify the relative extent of contemporary (2000 – 2022) logging and the condition of the remaining native forest and woodland (quantified by measuring the similarity of a current ecosystem to a historical reference state with high ecological integrity). Using these data, we measure the impacts on distinct vegetation types and on 484 terrestrial forest-dependent now-threatened species. We show that more than half (29 million ha) of pre-1750 (pre-European colonization of Australia) native forest and woodland vegetation in NSW has been lost. Of the remaining 25 million ha, 9 million ha is degraded. We found contemporary degradation from logging affected 244 forest-dependent now-threatened species that had already been affected by this historical deforestation and degradation, but the impacts varied across species and vegetation types. We found that 70 now-threatened species that were impacted by historical deforestation and degradation and continue to be impacted by logging, now have ≤50% of their pre-1750 extent remaining that is intact (with three species now having <20%). By quantifying the historical impacts of deforestation and degradation, our research sets the impact of contemporary degradation from logging in perspective and highlights shortfalls in current environmental assessments that fail to consider appropriate baselines when reporting on overall impact. Future land management decisions need to consider not only the extent of remaining habitat based on pre-1750 extents, but also its condition.

**Article impact statement:** The impact of logging needs to be placed in perspective by considering past losses and degradation due to human land use decisions.

## Introduction

It is estimated that the global native forest and woodland estate harbours up to 100 million different species (80% of all terrestrial plants and animals, (United Nations 2021)), sequesters a net 7.6 billion metric tonnes of CO2 per year (1.5 times more carbon than the United States emits annually) (Harris et al. 2021), and provides essential ecosystem services that directly support more than 1.6 billion people (UNEP and FOA 2020; IUCN 2021). Despite their critical role in helping humanity overcome the challenges of biodiversity loss, climate change, and achieving global sustainability (FAO and UNEP 2020), forests and woodlands are among the most structurally altered terrestrial biomes on Earth (Williams et al. 2020).

Deforestation (defined as the outright removal and permanent conversion of forest or woodland to a non-woody land use) and degradation (defined as the gradual process of function or biomass decline, changes to taxa composition, or erosion of soil quality (European Union Reducing Emissions from Deforestation and forest Degradation 2022) are both well-known urgent environmental problems, driving considerable global protection agendas (e.g., (Leaders pledge for Nature 2022; Secretariat of the Convention on Biological Diversity 2022) and restoration goals (The Bonn Challenge 2020; Trillion Trees 2021). Although deforestation and degradation rates are increasingly well documented at both global (Beyer et al. 2019; Grantham et al. 2020a; Williams et al. 2020) and national scales (Grantham et al. 2020b; Williams et al. 2021a), their cumulative impact on vegetation types and forest-dependent species’ habitat over the long timescales humans have been impacting forests, is often ignored in contemporary impact assessments that inform future land-use decision making.

In Australia, there have been published studies examining the extent of deforestation and fragmentation of forests between pre-European colonisation (1750) to 2009 (Bradshaw 2012) and the extent of recent deforestation of forest-dependent threatened species habitat (Ward et al. 2019). There has also been analysis on the drivers and trends of deforestation in Australia (Evans 2016), and individual species-specific assessments of threats which include deforestation and degradation (e.g., (Coxen’s fig-parrot recovery team 2005; Menkhorst 2008), yet there has been no quantification of the extent of both historical deforestation and degradation for specific Australian vegetation types or for forest-dependent threatened species. Furthermore, no assessment of the ongoing impact of one key, manageable contemporary driver of degradation, native forest and woodland logging, has been considered in this holistic context.

While the failure to place logging within a historical conservation context is commonplace, it means current environmental assessments and environmental accounting do not address shifting baselines, making them problematic to use when making decisions about future management (Papworth et al. 2009; Lindenmayer & Laurance 2012; Soga & Gaston 2018). It also impedes accurate reporting on how much species’ habitat has really been lost or degraded (Ward et al. 2022a). Placing the impacts of historical and contemporary land use in context when it comes to species and ecosystems will help establish how feasible it will be for nations to achieve international agreements such as the Sustainable Development Goals (United Nations Sustainable Development Goals 2015), the Kunming-Montreal Global Biodiversity Framework (Secretariat of the Convention on Biological Diversity 2022), and Leaders Pledge for Nature (Leaders pledge for Nature 2022) if deforestation and degradation continues. At local management scales, it will also allow for more complete assessments when analysing the likely consequences of current contemporary degradation through activities like logging, because when these activities are undertaken in isolation and not considered in long-term land management histories, relatively small areas that are logged (or planned to be logged) can be presented as inconsequential. This is especially true when small areas are presented in terms of the former historical areas without also acknowledging how much of that former area is already lost or degraded (Rittenhouse et al. 2010).

Here, using New South Wales (Australia) as a case study, we provide an assessment of historical forest and woodland deforestation since 1750 (pre-European colonization of Australia) – 2021 alongside an assessment of degradation (pre-European colonization of Australia – 2018). We then assess the impacts of logging (2000-2022), one of several ongoing contemporary drivers of degradation, against this historical deforestation and degradation to capture an overall assessment of how ongoing drivers are affecting vegetation types and now-listed terrestrial (and semi-terrestrial) forest-dependent now-threatened species. Australia has many endemic flora and fauna and is one of 17 mega-biodiverse. Many of these endemic species have suffered significant declines in recent decades, with Australia having 1,869 species listed as threatened with extinction by the Federal Government, with 104 taxa now extinct (Commonwealth of Australia 2022a). Deforestation and degradation is a major cause of this species crisis (Fischer & Lindenmayer 2007; Mac Nally et al. 2009; Ford 2011; Ward et al. 2021; Kearney et al. 2023), yet decisions about the management of forests and woodlands typically do not consider the impacts of contemporary actions in the context of the historical deforestation and degradation (Lindenmayer & Laurance 2012).

By integrating deforestation and degradation data with vegetation and species distribution maps, we showcase a method that enables assessments and reporting on the impacts of contemporary logging that also considers historical deforestation and degradation. Importantly, while we cannot change historical deforestation and degradation beyond focussing on targeted restoration where possible (Mappin et al. 2021), key stakeholders and decision-makers can prevent further degradation from logging, especially in areas that are clearly the last remaining intact habitats that are critical for securing threatened species (Ward et al. 2022). The information provided here allows for these decision makers in New South Wales to truly consider the overall impact of contemporary actions like logging in the context of the overall historic deforestation and degradation (Lindenmayer & Laurance 2012). We argue that these types of assessments should be used as the basis of future land management assessments when it comes to considering the impact of any planned activity that degrades or destroys intact vegetation.

## Methods

### Study region

Our study covers the state of New South Wales (NSW, Australia), proportionally, the second most forested and woody state of the Australian continent. NSW supports more than 1,600 plant community types (NSW Government 2022) and 532 now-threatened species (233 of which are endemic to NSW) listed as Vulnerable, Endangered, or Critically Endangered under the Environment Protection and Biodiversity Conservation (EPBC) Act 1999, Australia’s key federal piece of environmental legislation (Commonwealth of Australia 1999).

### Threatened species data

To examine the potential impact on now-threatened species, we first created a list of nationally listed threatened species, using the information available from the federal Government’s website (Commonwealth of Australia 2022a). The Maxent modelled distributions of each of these taxa were then sourced from a publicly available database provided by the Federal Government’s Department of the Environment and Energy’s (retrieved 15th October 2022). Maxent (or maximum entropy modelling) predicts species occurrences by considering the limits of environmental variables of known locations (Elith et al. 2011). The information used to create the Maxent modelled distributions of threatened species is sourced from a range of government, industry, and non-government organisations with expert opinion and reference to published information. As of 15^th^ October, 2022, there were 532 threatened species, subspecies, and populations (hereafter referred to as ‘taxa’) occurring in NSW (with at least 1% of total distribution), with most distributions generalised to ~1km^2^ or ~10km^2^ grid cells (Commonwealth of Australia 2022b). We refined this spatial data to only 484 forest-dependent terrestrial or peri-terrestrial species (where forest-dependent is defined as a taxon that intersects with ≥5% of forest or woodland vegetation groups mapped in the pre-1750 National Vegetation Information System (NVIS 6.0; (Commonwealth of Australia 2020)). Peri-terrestrial taxa included in this study included all threatened frogs. We do not include aquatic or some peri-terrestrial species such as fish, crayfish, or turtles as the impacts and method to measure impact is vastly different to terrestrial systems. We use now-threatened taxa as the focus of this study, recognising that historical deforestation and degradation impacts have most likely contributed to their contemporary threatened status.

### Historical clearing map

We identified the extent of deforestation from 1750-2021 using five lines of evidence. The first was the Australian Government’s National Vegetation Information System (NVIS 6.0; (Commonwealth of Australia 2020)). This spatial layer summarizes Australia’s present native vegetation, classified into 33 Major Vegetation Groups determined by structural and floristic information including dominant genus, growth form, height, and cover. NVIS also includes a category identified as ‘Cleared, non-native vegetation, buildings’ at 1 ha resolution. The second layer was the State Vegetation Type Map (SVTM), which is a regional-scale map using a vegetation classification hierarchy, including vegetation formations, vegetation classes, and plant community types (NSW Government 2022). The SVTM also identifies ‘non-native’ and ‘no vegetation’ used here to identify deforestation as of 2021. We used two forest and woodland loss spatial layers (Ward et al. 2019; Thomas et al. In prep) – derived from Australia’s National Carbon Accounting System (NCAS) forest and woodland cover dataset – to identify forest and woodland clearing from 2000-2021. The fifth layer utilised was the Habitat Condition Assessment System (HCAS) version 2.1 applicable to 2018 (Harwood et al. 2021; Williams et al. 2021a). HCAS is a new way of combining environmental data, remote sensing data, and intact condition reference sites to provide a consistent estimate of habitat ‘condition’ or quality for all locations across Australia (Harwood et al. 2016). Condition is defined as the predicted capacity to support the wildlife expected in a given area under natural conditions (Williams et al. 2021a). HCAS v2.1 provides, for every 250m^2^ pixel, a score from 0 – 1, which can be broken up into five ordinal categories including residual (0.81 – 1), modified (0.61 – 0.80), transformed (0.41 – 0.60), replaced (0.21 – 0.40), and removed (0.20 – 0), to approximate the generalised states and transitions narrative proposed by Thackway and Lesslie 2008 for Australian native vegetation, and as used in National state of the environment reporting (Thackway & Lesslie 2008; Williams et al. 2021b). To create one deforestation map from 1750 – 2021, we combined the NVIS ‘Cleared, non-native vegetation, buildings’ data, SVTM ‘non-native’ and ‘no vegetation’ data, the two forest and woodland loss datasets, and HCAS ‘replaced’ and ‘removed’ data. To identify just deforestation of forests and woodlands, we overlayed this with forest and woodland vegetation groups as per NVIS 1750.

We overlayed the historical forest and woodland clearing map with individual taxa distribution maps to quantify impact for the 484 EPBC Act listed terrestrial, forest-dependent now-threatened taxa within NSW. To quantify how much of the remaining taxa distribution was degraded, we then overlayed the modified (0.61 – 0.80) and transformed (0.41 – 0.60) pixels from the HCAS map with individual taxa distribution maps. We acknowledge that in some cases, calculations of loss and degradation for specific taxa may be an underestimate as we use only distributions for where species occur currently, and not their full historical distribution. However, for most taxa, their pre-European colonisation distributions are unknown as vegetation was destroyed or degraded before Western documentation occurred. Where historical distributions are larger than the current estimates, our analysis will underestimate the extent of historical impacts of deforestation and degradation. To ensure no double counting of deforestation or degradation, we removed any logging overlaps (described below) with the historical deforestation layer and the historical degradation layer. Note, when reporting on species impacts, we considered only the forest or woodland portion of habitat that is within NSW.

### New South Wales logging data

Logging data were retrieved from Forestry Corporation of New South Wales’ (FCNSW) Open Data Site (FCNSW 2021) (retrieved 17th November 2022). These data show forests and woodlands that have been logged (using selective logging techniques) between 2000 – August 2022. We removed all plantations (provided by FCNSW in August 2021) from the logging layer because plantations were captured in the above deforestation layer. We also removed any logged area from the degradation layer described above to ensure no double counting across the spatial layers. We removed any areas that were logged prior to 2000 to align with the inauguration of the EPBC Act and listed threatened species. We intersected taxa distributions to the boundaries of completed logging to estimate the overall extent of degradation impacts by logging. To assess the condition of remaining forest and woodland within taxa distributions (i.e., not deforested, degraded, or logged), we intersected all taxa distributions with the residual (0.81 – 1), modified (0.61 – 0.80), and transformed (0.41 – 0.60) pixels in HCAS. Here, we refer to residual as ‘intact’, and modified and transformed is described as ‘degraded’.

Under the EPBC Act, a taxon may be listed as Critically Endangered, Endangered, or Vulnerable for many reasons, including if it experiences a population size reduction of >80%, >50%, or >30%, respectively, measured over the longer of ten years or three generations (where threats are ongoing and unresolved, and given temporal considerations). A decline in area of occupancy, extent of occurrence, and/or quality of habitat exceeding these thresholds can be taken as an indicator of equivalent population declines. Here, we assessed the combination of historical deforestation and degradation, contemporary logging, and remaining condition of forest and woodland within taxa distributions against such criteria.

## Results

By 2021, the total forest and woodland remaining in NSW was ~25 million ha. The amount of forest and woodland lost due to deforestation is approximately 29 million ha or 54% of the 1750 native forest estate (55 million ha) (**Fig. 1**). Most deforestation has been concentrated along the east coast of NSW, within several major vegetation groups including Eucalypt Woodlands (10 million ha (35%) remaining), Eucalypt Open Forests (5 million ha (48%) remaining), and Eucalypt Open Woodlands (1 million ha (35%) remaining), based on the National Vegetation Information System.

**Figure 1.**
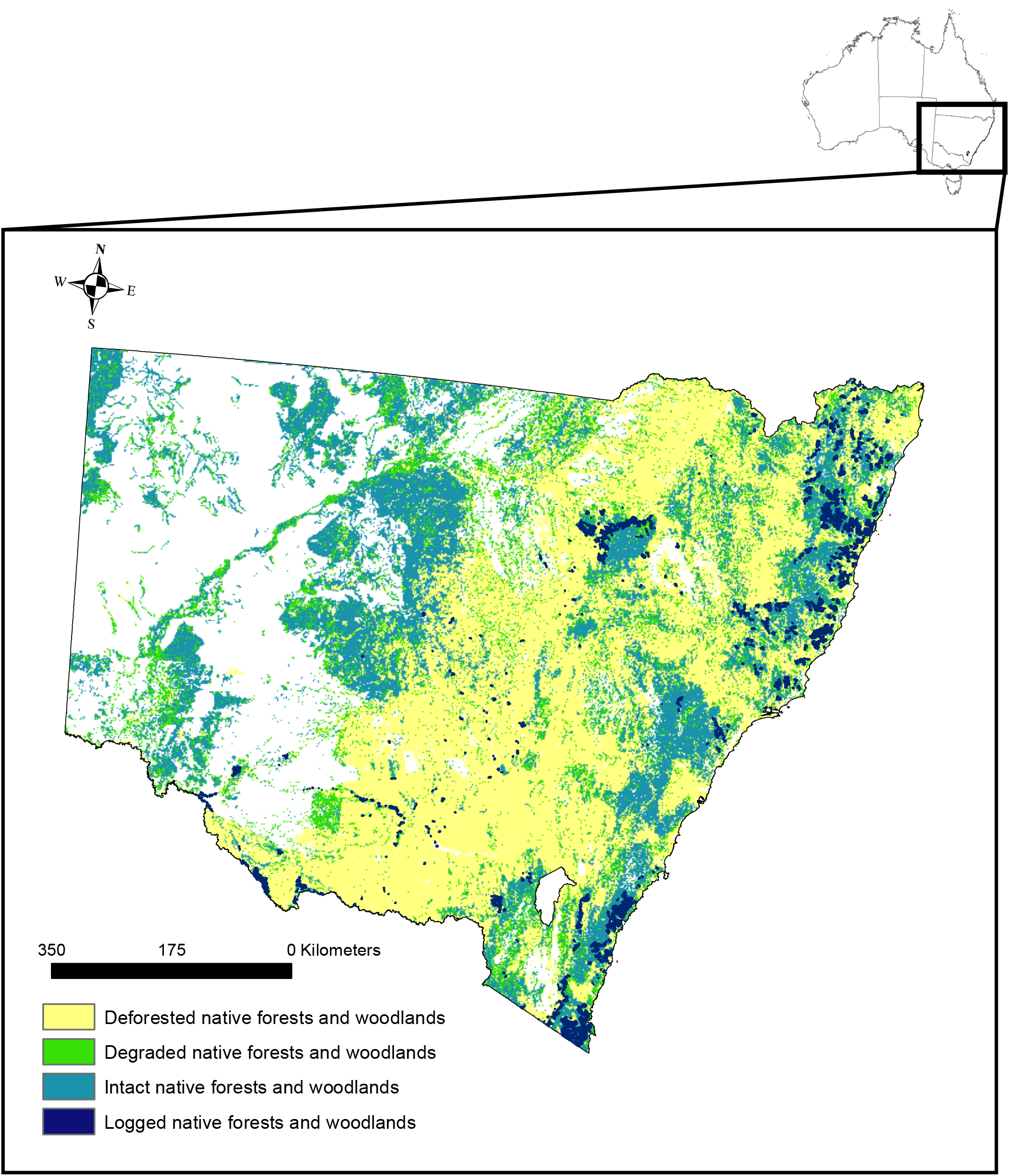
Map of 2000 – 2022 logged areas in NSW represented in dark blue with dark blue border for increased visibility at this scale. From 1750-2021, cleared forests and woodlands is represented in yellow, while remaining degraded native forests and woodlands is represented in green. Remaining intact native forests and woodlands is represented as teal.

Of the remaining forests and woodlands, approximately 16 million ha (30% of all pre-European forest and woodland) is intact, and 9 million ha is degraded. While some vegetation groups have not been heavily cleared, they have been heavily degraded. When assessing the condition of remaining forest and woodland vegetation groups, 72% of remaining Casuarina forests and woodlands is degraded, 45% of remaining Melaleuca forests and woodlands is degraded, and 39% of remaining Eucalypt open woodland is degraded.

We found that contemporary degradation in the form of logging continues in 12 (out of 15) of NSW’s major forest and woodland vegetation groups. Our analysis found the area of logged forests constitutes mostly Eucalypt tall open forests (150,000 ha) and Eucalypt open forests (135,000 ha). The total extent of logging within NSW from January 2000 - August 2022 was 435,000 ha.

### Potential impact of deforestation and degradation on now-threatened taxa

All 29 million ha of historical deforestation overlapped with the distributions of at least one of the 484 now-threatened taxa considered in this study. In total, 459 now-threatened taxa have potentially been impacted (>1% of forest or woodland distributions) by historical deforestation. The extent of overlaps between deforestation and species distributions ranged from 1% - 99% (mean = 35%, median = 32%). Flora that have the lowest proportional distribution remaining include *Homoranthus bruhlii* (1% of woody distribution remaining), *Hibbertia puberula subsp. Glabrescens* (1% of woody distribution remaining), and *Eucalyptus alligatrix subsp. miscella* (4% of forested distribution remaining). All three species occur only in NSW. Fauna that have the lowest proportional distribution remaining include Sloane’s froglet (*Crinia sloanei*; 3% of woody distribution remaining in NSW), Key’s matchstick grasshopper (*Keyacris scurra*; 13% of woody distribution remaining in NSW), and golden sun moth (*Synemon plana*; 14% of woody distribution remaining in NSW).

Our results show that all areas logged between 2000 and 2022 overlapped with the distributions of at least one federally listed now-threatened taxa and in total, 244 taxa (50% of species assessed) were potentially impacted by logging (Appendix S1). Of these 244 taxa, 25 are listed as Critically Endangered, 79 as Endangered, and 140 as Vulnerable. The flora with the highest proportion of NSW forest and woodland distribution overlapping with logging includes stiff groundsel (*Senecio behrianus*; 79% of remaining woody distribution overlapping with logging), floodplain Rustyhood (*Pterostylis cheraphila*) (75% of remaining woody distribution overlapping with logging), and Narrabarba wattle (*Acacia constablei*, 32% of remaining woody distribution overlapping with logging; **Fig. 2**). The fauna taxa with the highest proportion of NSW distribution overlapping with logging include long-footed potoroo (*Potorous longipes*, 14% of remaining woody distribution overlapped with logging), long-nosed potoroo (*Potorous tridactylus trisulcatus*, 12% of remaining woody distribution overlapped with logging) and southern brown bandicoot (*Isoodon obesulus obesulus*, 9% of remaining woody distribution overlapped with logging; Table S2). Taxa with the most distribution by area overlapping with logging include koala (*Phascolarctos cinereus*, 400,000 ha), south-eastern glossy black-cockatoo (*Calyptorhynchus lathami lathami*, 370,000 ha), and Australian painted snipe (*Rostratula australis* (SE mainland population), 330,000 ha).

**Figure 2.**
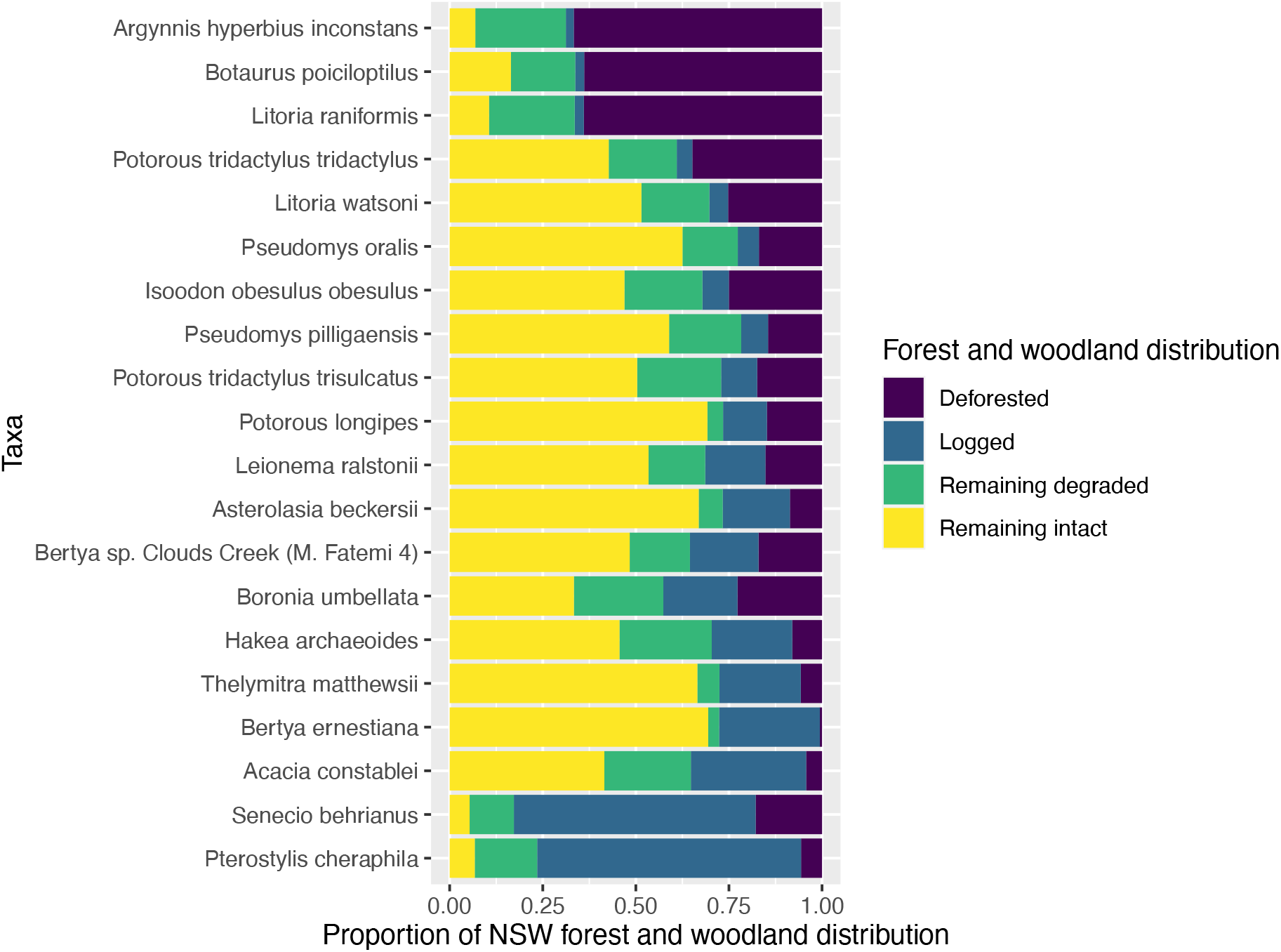
Proportional impact of historical deforestation (purple), logging (blue), remaining degraded (green), and remaining intact (yellow) highlighting the top 10 flora and top 10 fauna likely impacted by logging.

**Figure 3.**
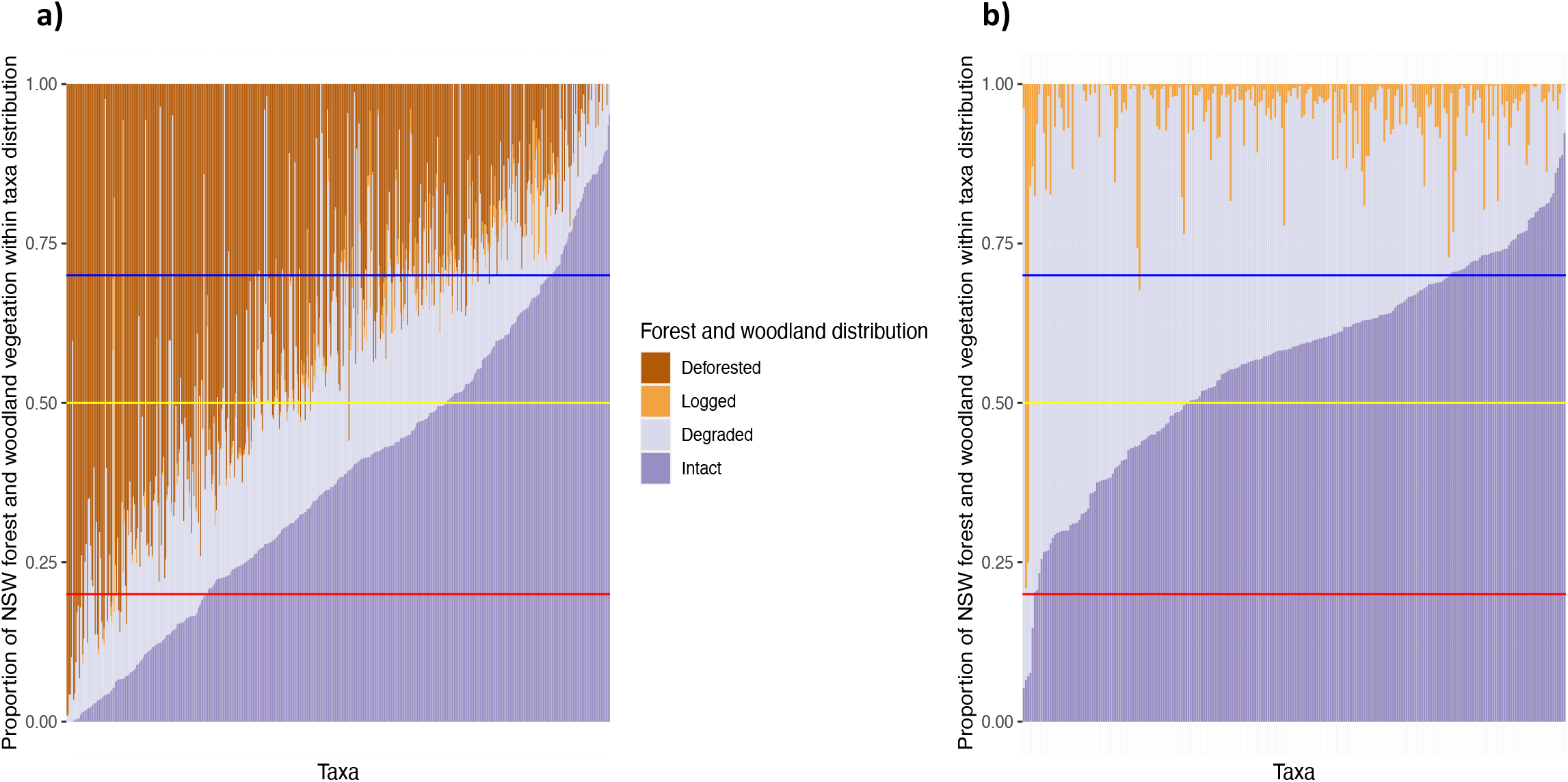
**a)** Proportion of each taxa’s distribution (x-axis) impacted by both historical deforestation, historical degradation, and contemporary logging. We show the combination of forest and woodland distribution potentially impacted from deforestation (brown) and logging (yellow), and highlight the condition of the remaining forest and woodland vegetation within taxon distributions as either degraded (light purple) or intact (dark purple). **b)** Proportion of each taxa’s distribution (x-axis) impacted by contemporary logging (yellow) only, highlighting the condition of the remaining forest and woodland vegetation within taxon distributions as either degraded (light purple) or intact (dark purple). The horizontal lines on both figures indicate EPBC Act threat status thresholds of ≤70% (yellow), ≤50% (blue) and ≤20% (red) for Vulnerable, Endangered and Critically Endangered, respectively. Both figures are ordered based on proportion on forest and woodland distribution remaining that is intact.

We assessed the condition of the remaining forest and woodland distribution after historical deforestation and degradation, and contemporary logging against EPBC Act listing criteria (see methods) and found that three taxa have ≤20% intact forest and woodland distributions remaining. These were the Endangered Sloane’s froglet, of which 5% of remaining forest and woodland distribution is intact, the Vulnerable Glenugie karaka (*Corynocarpus rupestris subsp. rupestris*), of which 9% of remaining forest and woodland distribution is intact, and the Vulnerable Mueller daisy (*Brachyscome muelleroides*), of which 17% of remaining forest and woodland distribution is intact. The distributions of all three species continue to be impacted by logging. Sixty-seven taxa have between 20 - ≤50% of remaining forest and woodland habitat intact (six are Critically Endangered, 24 are Endangered, and 37 are Vulnerable) and 102 taxa have between 50 - ≤70% of remaining forest and woodland habitat intact (11 are Critically Endangered, 29 are Endangered, and 62 are Vulnerable).

## Discussion

Total deforestation of native forest and woodland in NSW from 1750 - 2021 was ~29 million ha, equating to 54% of all native forests and woodlands across the state or an area approximately the size of New Zealand. Deforestation reduced the distributions of many native species, including 459 now federally listed forest-dependent now-threatened taxa we assessed here. Despite these large historical impacts, the habitat of 244 of these now-threatened taxa continues to be logged. We found that 70 threatened species that were impacted by historical deforestation and degradation and continue to be impacted by contemporary logging now have ≤50% of their pre-European extent remaining that is intact. Two of these species (Sloane’s froglet and Mueller daisy) have <10% of their NSW forest and woodland distribution remaining.

While deforestation has clear, immediate impacts on biodiversity such as removing habitat, resources, food, and shelter, forest degradation is more subtle and often overlooked (Thorn et al. 2020). Degradation, such as logging, is the gradual process of forest biomass decline, changes to taxa composition, or erosion of soil quality (European Union Reducing Emissions from Deforestation and Forest Degradation 2022). Native forest logging has severe degrading impacts on forests and subsequent forest-dependent biodiversity, such as reduction of critical resources necessary for taxa survival, including food, shelter, and breeding areas (Lindenmayer et al. 2013; Ashman et al. 2021). Another significant impact from logging is the network of roads needed to carry timber out of forests and woodlands and into processing facilities such as sawmills. Road networks cause major problems such as facilitating invasive predator access (e.g., cats (*Felis catus*), dogs (*Canis lupus familiaris*), and foxes (*Vulpes vulpes*)), and the spread of pathogens (e.g., chytrid fungus (*Batrachochytrium dendrobatidis*) and *phytophthora cinnamomi*; (Boston 2016) (Boston 2016). These permanent drivers of degradation often have larger effects than the reduction of vegetation from constructing the road. Some of these impacts may not manifest for several years or decades after logging, but often this extinction debt must be paid (Szabo et al. 2011).

Logging native forests can lead to increases in the severity and frequency of wildfires (Taylor et al. 2014; Lindenmayer et al. 2020). Logged areas burned with significantly increased severity during the record 2019/20 fire season in NSW (Lindenmayer et al. 2022). Fire is a critically important ecological disturbance that drives landscape heterogeneity, recruitment, community composition, and ecosystem function (Koltz et al. 2018; McLauchlan et al. 2020). For example, many obligate seeding plant taxa which are killed by fire require inter-fire intervals that are long enough to allow recruiting individuals to reach maturity and set seed into the seedbank to ensure their continued persistence (Keith 1996). However, drivers such as anthropogenic climate change and logging are resulting in fire becoming more prevalent, larger in scale, and occurring outside of historical fire seasons (Dowdy et al. 2019; Lindenmayer et al. 2020). This not only causes direct mortality of taxa, removal of habitat, and reduction in resources, fires can also shift ecosystems to different states, or interact with existing threatening process resulting in further degradation (Suding et al. 2004). Such changes in fire regimes represent a key threatening process to more than 800 Australian threatened native species, and 65 threatened communities (DAWE 2020; Ward et al. 2020). While we cannot control future wildfires, we can implement actions that will help reduce their impact, such as decarbonization and ending native forest logging.

In addition to destruction from deforestation and ongoing contemporary degradation from logging, many of these same species must survive other drivers of degradation such as invasive taxa (Legge et al. 2017), disease (Fensham et al. 2020), altered water regimes (Lintermans et al. 2020), climate change (Graham et al. 2019), pollution (Richman et al. 2015), and overexploitation (Ward et al. 2021). Our findings on the impacts of deforestation and degradation are critical for assessing the impact upon taxa, as currently proposed destructive actions are individually assessed and do not consider the cumulative impact (Dales 2011; Tulloch et al. 2016a) or are not assessed at all (Ward et al. 2019). In addition, under both federal and state legislation, actions are assessed without any historical context. While small amounts of impact each year may seem insignificant; the combined deforestation and degradation of habitat over 250 years can lead to taxa extinction via many small modifications of habitat (i.e., “death by a thousand cuts”) (U.S. Environmental Protection Agency 1999; Tulloch et al. 2016b; Reside et al. 2019).

One clear finding in this study is that it seems the state of New South Wales is locking in extinction through legislative inadequacies. Between 2000 - 2022, NSW logged approximately 435,000 ha of native forest and woodland (all of which overlaps with the distributions of at least one now-threatened forest-dependent taxa). This impact assessment is likely an underestimate given the rate of new discoveries of species in Australia. In case and point, in 2022 alone, scientists discovered an additional 139 new species in Australia (CSIRO 2022). While the EPBC Act is the primary legislation aimed at protecting biodiversity, the Regional Forest Agreements (RFA) Act 2002 is geared towards ensuring access to forests whilst ensuring the conservation of forest biodiversity and protection of environmental integrity (The Department of Agriculture water and the environment 2020). Unfortunately, RFAs have been exempt from following EPBC Act protections (Lindenmayer & Burnett 2022), even when there are clear breaches such as degrading threatened taxa habitat which is likely to have a significant impact on those species (Commonwealth of Australia 1999). Therefore, while logging in NSW has degraded 435,000 ha of native forest across 244 threatened taxa distributions, it is still legal under current legislation (Ashman & Ward 2022).

We do note that Australia (alongside many countries) have recently signed international agreements to halt taxa extinctions (e.g., Convention of Biological Diversity), prevent further forest degradation (Glasgow Climate Pact, Glasgow Leaders’ Declaration on Forests and Land Use), and reverse biodiversity loss (e.g., Natures Pledge). The NSW Government has also made commitments to enhance nature conservation including stopping extinctions inside protected areas, stabilising, or improving the trajectory of all threatened taxa, and removing threatened taxa from the threatened species list (NSW Government 2021a). Notably, the NSW Government’s Koala Strategy commits to doubling koala numbers by 2050 (NSW Government 2021b). Many countries and jurisdictions are now legislating that commodity production (such as beef, cocoa, soy, and timber) must not contribute to deforestation and degradation. For example, the European Union passed new laws December 2022 to ensure there is now a due diligence process to demonstrate that imported produce have not contributed to deforestation or degradation (European Commission 2022). In addition to driving species to extinction, and possible inaccessibility to markets, it is also not economically viable. A recent analysis measured the stream of costs and benefits over a 30-year period to 2051 and found that ending native forest logging in the southern areas of NSW and using these areas for their environmental and recreational benefits, could result in a $61.96 million saving of taxpayer money (Frontier Economics & Macintosh 2021). Unfortunately, there has been no commitment by the NSW Government to end logging in native forests. To ensure Australia meets its international, national, and state commitments on biodiversity conservation, sufficient habitat needs to be protected, managed, and/or restored to support viable populations of target taxa.

### Caveats and limitations

We recognize that our historical impact analysis of forest and woodland within the distributions of threatened taxa is likely an underestimate as we have used known and likely to occur distributions based off recent records of taxa. This does not include the contractions of the historical distribution that have occurred due to habitat loss and other perverse threatening processes such as invasive species and changed fire regimes (Ward et al. 2022). There are also many non-threatened species, such as aquatic and peri-terrestrial species, that have experienced huge historical habitat loss and ongoing contemporary degradation that have not been captured here. Unfortunately, the NSW Forestry Corporation do not report their impacts on threatened taxa, so we were unable to compare our results with existing data captured by industry.

## Conclusion

Despite strong evidence that Australia’s biodiversity is suffering major declines, degradation by multiple sources including logging, continues. Policy makers and the community must recognize and account for the critical values of intact native forests such as high biodiversity, mitigating climate change, services to people (such as clean water provision and air purification), and low fire severity. This research showcases a way to measure the historical impacts of deforestation and degradation, as well as contemporary degradation from logging. We highlight how threatened taxa may be impacted and emphasize the critical importance of considering these long-term impacts in contemporary settings when it comes to forest management in New South Wales. These holistic, contextual assessments need to be the basis of all future land management decisions.

## Acknowledgements

We acknowledge the Traditional Owners of Country throughout New South Wales, we recognise their continuing connection to land, waters and community and acknowledge that Traditional Owner sovereignty was never ceded. We pay our respects to their cultures and elders past, present, and emerging.

